# Mapping social profiles in childhood and adolescence: associations with cognition and brain structure

**DOI:** 10.64898/2026.04.20.719698

**Authors:** Estherina Trachtenberg, Alexa Mousley, Maria Jelen, Duncan Astle

## Abstract

**Objective:** Social difficulties are transdiagnostic in childhood, but their heterogeneity is poorly characterised and rarely treated as a primary neurodevelopmental phenotype. This matters because childhood and adolescence are sensitive periods for peer relationships and brain development. We used data-driven modelling and non-linear mapping to derive social profiles and test their clinical, cognitive, and neural correlates.

**Methods:** Participants were 992 children aged 5–18 years from CALM (Mage = 9.6). Social items from the SDQ, CCC-2, and Conners-3 were modelled using a regularised partial correlation network to derive core social dimensions. A self-organising map captured graded social profiles. Simulated archetypes, SVM-based island identification, and permutation testing defined profile regions and centroid-distance scores. Profiles were related to referral, diagnosis, cognition, BRIEF indices, and T1-derived MIND network structure in an MRI subsample (n=431).

**Results:** We identified four profiles: social engagement, friendship difficulties, social withdrawal, and peer victimisation. Profile expression tracked variation in referral and diagnostic pathways. Social withdrawal showed the clearest disadvantage across cognitive domains, whereas social engagement was associated with fewer executive function difficulties across BRIEF indices. MIND strength components covaried with profile expression (a significant PLS latent variable, p = 0.02), with covariance strongest for social withdrawal and peer victimisation.

**Conclusions:** Childhood social functioning organises graded signatures that relate to clinically relevant pathways, cognitive and executive outcomes, and brain structure. Profiling social signatures provides a scalable framework for identifying social need beyond diagnostic categories, motivating studies to test directionality and improve developmental outcomes.

## Introduction

Childhood and adolescence are critical periods in which social relationships become central to everyday functioning and long-term development. These years coincide with ongoing maturation of prefrontal and distributed control systems that support executive function, learning, and emotion regulation^1,2^. During these years, peer relationships are not only a context for companionship, they are also a major arena for learning, emotion regulation, and establishing a sense of social safety^3^. Peer victimisation, in particular, is consistently associated with psychosocial maladjustment, later mental health difficulties, and lower academic achievement^4,5^. In contrast, prosocial behaviour and supportive peer relationships are associated with better academic outcomes and school engagement, and with fewer internalising and externalising symptoms^6–8^. Alongside observable peer behaviours and experiences, loneliness often co-occurs with peer difficulties, but it reflects a subjective appraisal of belonging and relationship quality that does not map perfectly onto peer-rated social functioning or social status^9,10^. Put simply, some children feel lonely despite appearing socially connected, while others have clear peer difficulties without feeling lonely.

In adults, loneliness and other social factors are linked to poorer cognition, whereas evidence in children and adolescents remains limited and focused mainly on mental health rather than cognitive development^11,12^. This gap matters because childhood and adolescence are sensitive, prolonged periods for developing executive functions and learning-related skills during ongoing maturation of prefrontal and distributed control systems^1,2^. During this window, social experiences are a salient developmental exposure. Friendships, peer belonging, and loneliness can shape daily opportunities for learning, support, and engagement with school, with consequences that may be amplified when developmental systems are still maturing^8^. Loneliness shows moderate stability in individual differences across development, and children who are lonelier than peers often remain relatively lonelier later, including into adulthood ^13,14^. This fits cascade accounts in which early peer difficulties heighten social threat and withdrawal, reducing opportunities for supportive friendships and sustaining loneliness over time^13^. Yet most work in children and adolescents has prioritised mental health outcomes, with far less mechanistic and longitudinal research testing how these social exposures relate to developing cognition and executive function.

Developmental stress frameworks provide one mechanistic account of how peer-based adversity could shape neurocognitive development^15^. Chronic exclusion and conflict may influence executive function and emotion regulation via sustained stress physiology and downstream corticolimbic alterations^10,16,17^. This framing motivates mapping heterogeneity in children’s peer relationship patterns to identify social risk signatures that co-occur with cognitive vulnerabilities, and to inform later longitudinal and intervention work that can test directionality.

A central challenge is directionality. Social withdrawal, rejection, bullying, and loneliness may shape cognitive and emotional development, while cognitive and regulatory difficulties may also increase vulnerability to peer difficulties, creating reinforcing cycles over time. Executive function and peer problems show similar bidirectional, developmentally embedded links^18^. Together, this supports treating social difficulties as a core developmental domain warranting direct characterisation and monitoring.

A practical implication is whether early identification of social difficulties can help flag children at risk for broader functional challenges, including cognitive and executive difficulties. A second implication is whether interventions that strengthen peer contexts, for example by increasing prosocial engagement and supportive peer interaction, improve learning trajectories rather than only reduce distress. Universal school-based social and emotional learning programmes show average gains in academic achievement alongside social–emotional outcomes^19^. Peer-learning approaches can support learning relative to working alone^20^, yet very few studies include executive function as an outcome.

In the current study, we map heterogeneity in social behaviour and peer relationships in the transdiagnostic CALM cohort^21^. Using parent-report items spanning peer relationships, social discomfort, prosocial behaviour, and peer adversity, we first estimate the empirical item-level structure to derive core social dimensions, then use a self-organising map to project child profiles onto a two-dimensional grid and capture non-linear patterns of co-occurrence^22^. Our aims are to (i) identify distinct, interpretable social profiles, and (ii) test associations with clinically relevant characteristics, including demographics, cognitive performance, and brain structure derived from T1-based morphometric inverse divergence . Although these analyses are cross-sectional and do not address directionality, they identify which social profiles show the strongest co-occurrence with neurocognitive variation, providing a foundation for future longitudinal and intervention studies that test developmental pathways (Figure 1).

**Figure 1.**
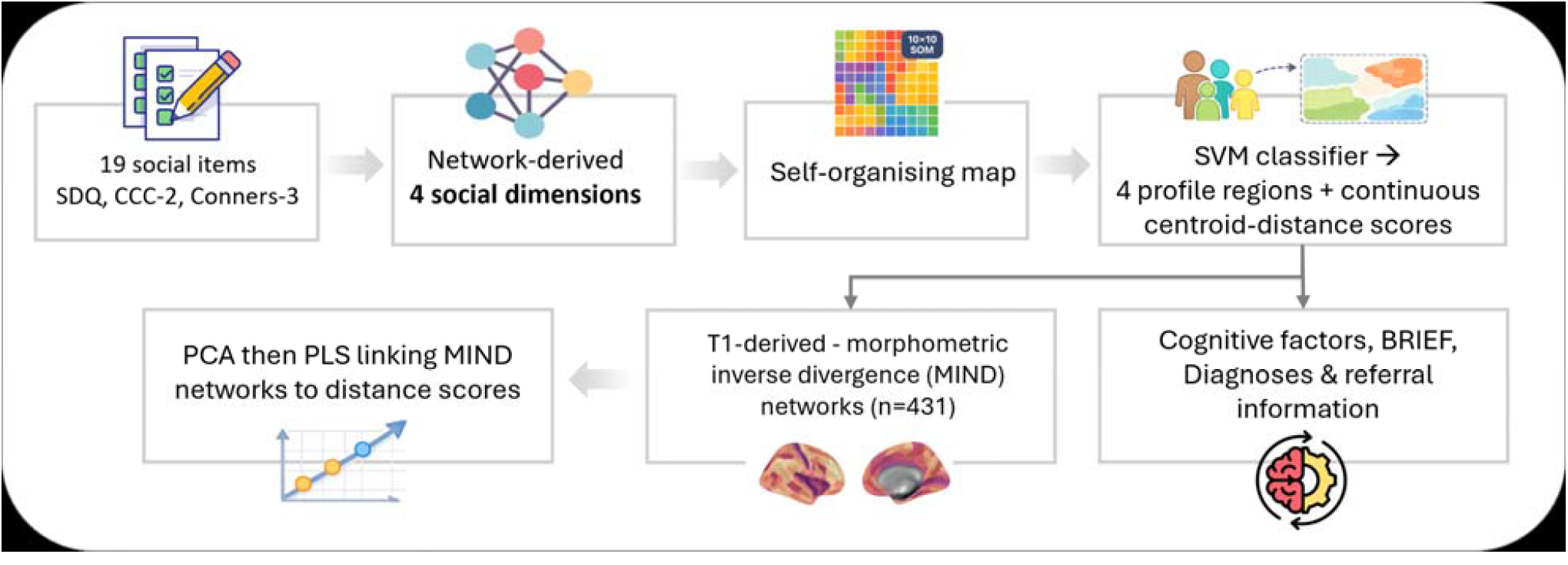
Overview of the analysis pipeline. Parent-report social items (SDQ, CCC-2, Conners-3; 19 items) were modelled using a regularised partial correlation network to estimate conditional associations among items and derive four core social dimensions. These dimensions were then used as input into a self-organising map to capture graded, non-linear configurations across individuals and to visualise the resulting social landscape. An SVM classifier was used to define four profile regions (“islands”) on the map, and continuous centroid-distance scores were computed to index each child’s degree of profile expression. Profile measures were then associated with referral and diagnostic information, cognitive factors, and BRIEF indices. In an MRI subsample (n = 431), T1-derived MIND networks were reduced with PCA to summarise major axes of morphometric network variation, and these components were related to social profile expression using PLS to identify multivariate brain–behaviour covariation patterns.

## Methods

### Participants

Participants were drawn from the Centre for Attention, Learning and Memory (CALM) cohort. The present study included 992 children (Mage = 9.6 years, SDage = 2.4, range 5 to 18), including 653 boys (65.8%) and 339 girls (34.2%) (Table 1). Neuroimaging analyses were conducted in a subsample of 431 children with associated MRI data (65.2% boys, 34.8% girls). Ethical approval: REC 13/EE/0157; IRAS 127675, full study procedures are described in the CALM protocol^21^. Children in CALM were referred by education or health professionals for difficulties with attention, learning, and/or memory, irrespective of diagnostic status. The sample included 805 referred and 187 non-referred children, with the latter recruited from the same schools and communities as a comparison group. Further details on diagnostic classifications and the list of diagnoses are provided in the Supplementary Materials.

**Table 1.**
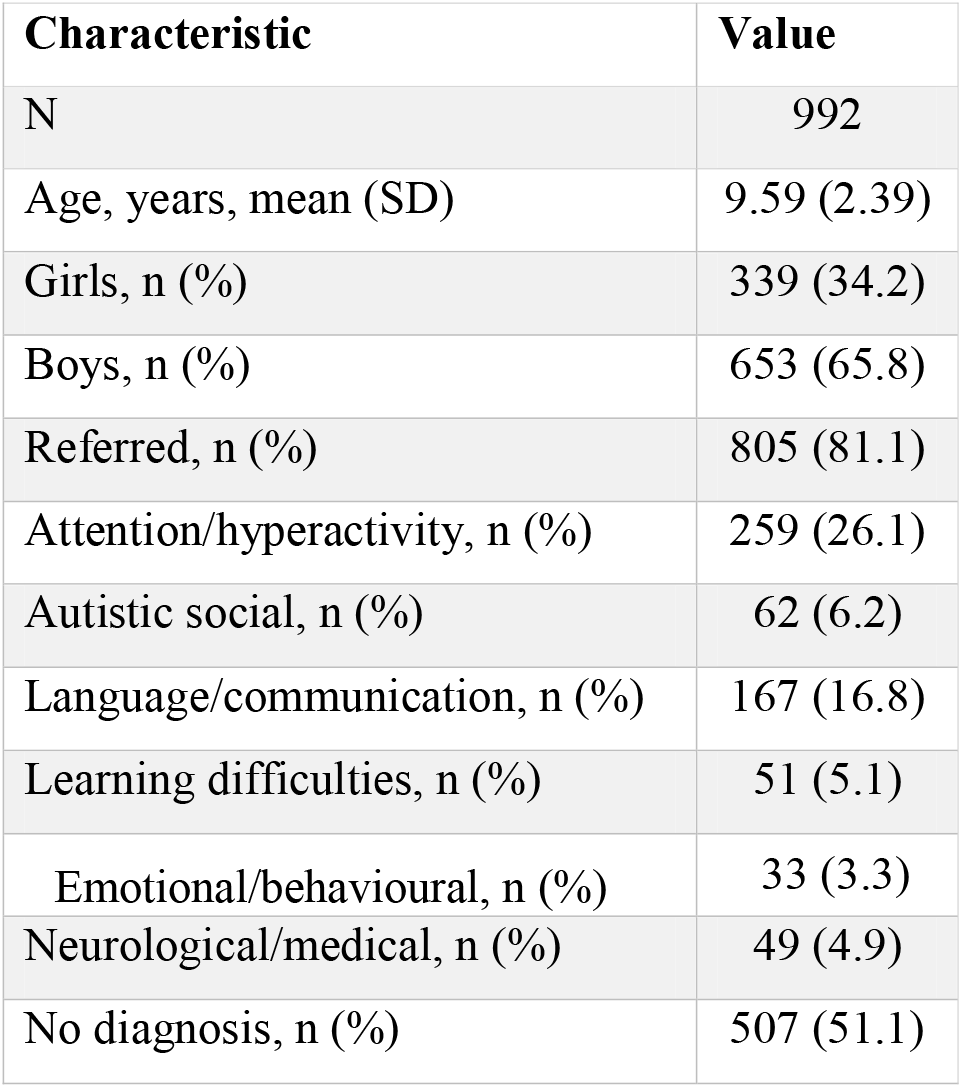
Sample characteristics.

### An overview of the analysis pipeline is provided in Figure 1

#### Social measures and network analysis

Social behaviour and peer relationship items were drawn from three parent-report questionnaires collected within CALM^21^: the SDQ, CCC-2, and Conners-3. Twenty-two items indexing social functioning were selected for analysis (Table S2). Item-level relationships were modelled using a regularised partial correlation network (EBICglasso), and communities were identified using Louvain modularity optimisation to derive social dimensions. One low-reliability community was excluded, leaving four retained dimensions (Text S1). Dimension scores were computed as the mean of z-scored items within each community and rescaled to the 0-1 range for SOM training.

### Self-organising map and profile classification

A self-organising map (SOM)^22^ was used to represent multivariate variation across children’s four social dimension scores in a two-dimensional space while preserving local neighbourhood structure^23^. The SOM was trained using a 10×10 rectangular grid with Euclidean activation distance in MiniSom. Training was run for 5,000 iterations with random initialisation, a Gaussian neighbourhood function (sigma=2) and learning rate 0.5, after which each participant was mapped to their best matching unit (BMU)^22^, defined as the node whose weight vector was closest to the participant’s input vector. Map quality was summarised using quantisation error and topographic error.

To identify interpretable regions within this graded space, we used a supervised classification approach anchored by simulated archetype profiles rather than standard clustering. Four archetypes were generated to reflect social engagement, friendship difficulties, social withdrawal, and peer victimisation. For each archetype, the corresponding dimension was set to the 95th percentile, and the remaining dimensions to the 50th percentile. A null archetype reflecting median values across all dimensions was included as a reference, and 500 simulated observations were generated for each archetype.

One-vs-rest support vector machine (SVM) classifiers with a radial basis function kernel^24^ were trained on the simulated archetype observations and applied to SOM neuron weight vectors. Each neuron was assigned to the archetype with the highest decision score. We then used a permutation-based island-size procedure to test whether same-label regions formed spatially contiguous islands larger than expected by chance. Islands were defined using 8-neighbour connectivity, and null island-size distributions were generated from 1,000 permutations in which SOM neuron weight vectors were randomly reassigned to grid locations and reclassified. For each archetype, the minimum island size required for significance was defined as the 99th percentile of its null distribution (p<.01). Significant islands were retained, and island centroids were computed as the mean row and column coordinates of neurons within each island.

Profile outputs were derived in two ways. For categorical assignment, children were assigned to the profile island containing their BMU, and those whose BMU fell outside all significant islands were classified as unassigned. For continuous profile expression, Euclidean distance was computed between each child’s BMU coordinate and the centroid of each significant island. Lower values indicated greater proximity to, and stronger expression of, the corresponding profile. Each child, therefore, had both a categorical profile label and four continuous centroid-distance scores (Figure 2B). Additional methodological details are provided in the Supplementary Materials.

**Figure 2.**
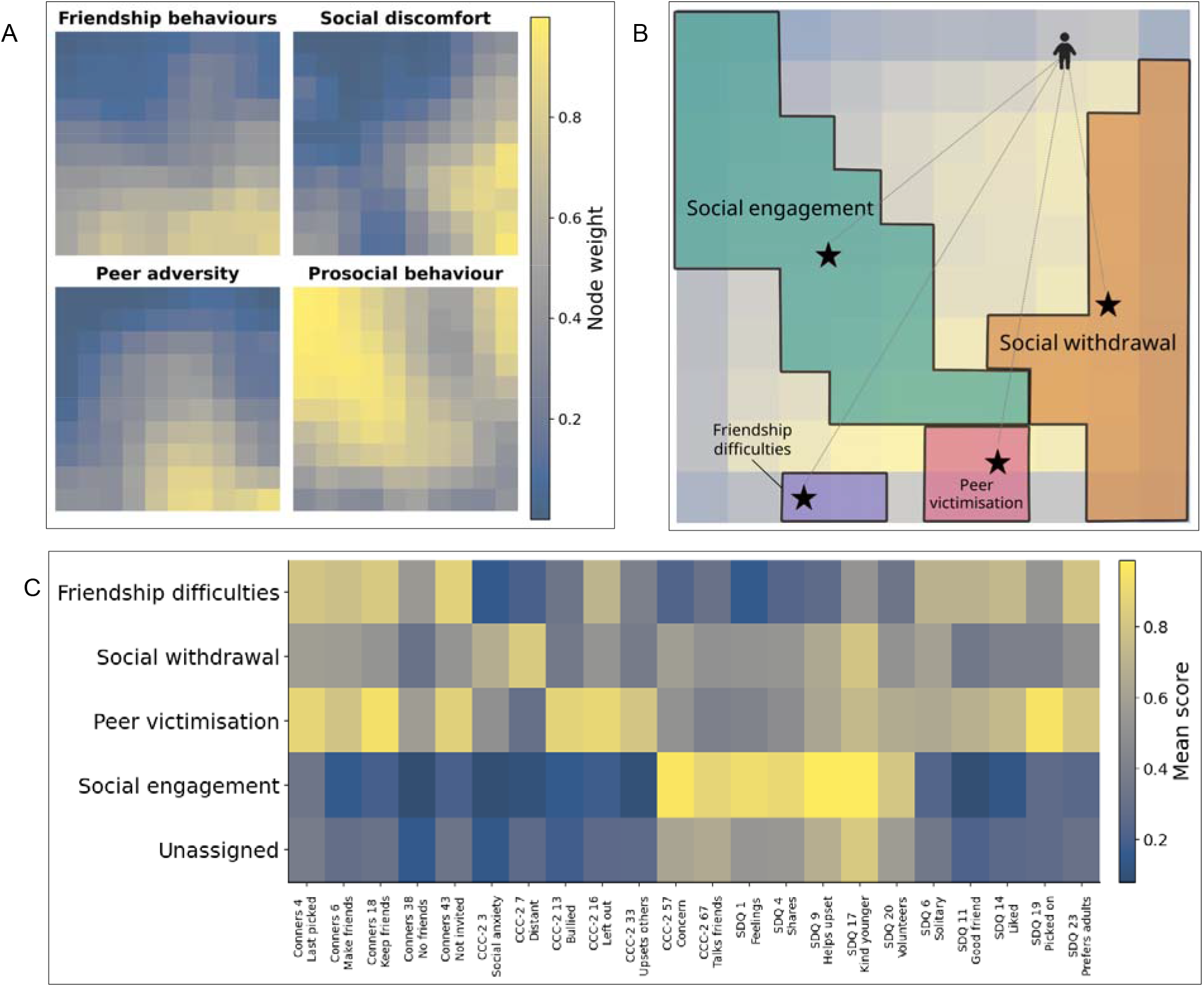
Social dimensions and profile islands on the SOM. (A) Component planes. Each SOM node stores a four-value weight vector, with one value for each social dimension used to train the map. Component planes plot these values one dimension at a time, showing how each dimension varies across the grid. Colours indicate the relative weight of each dimension at each node, with warmer colours representing higher values and cooler colours representing lower values. (B) Classification-based profile islands overlaid on the SOM U-matrix (distance map). Islands were identified using simulated archetypes and one-vs-rest SVM classifiers, and retained using a permutation-based island-size test (99th percentile of null island sizes, p < .01). Stars mark island centroids used for categorical membership (BMU within an island) and centroid-distance scores (distance to each centroid). Each child had one categorical profile label and four continuous centroid-distance scores, one for each profile. Dashed lines illustrate distances from an individual child to each centroid. (C) Per-item mean scores across the 22 social items by profile (including the unassigned group), illustrating item-level patterns characterising each profile.

### Cognitive measures and executive function

Cognitive performance was assessed using standardised measures of phonological processing, working memory, reasoning, vocabulary, and academic attainment^21^. Five cognitive summary measures were derived from age-standardised scores and rescaled to a 0-1 range, with higher values indicating better performance, for profile comparisons. Everyday executive function was assessed using the parent-report Behaviour Rating Inventory of Executive Function (BRIEF)^25^. Analyses focused on the three composite indices: the Global Executive Composite (GEC), Metacognition Index (MI), and Behaviour Regulation Index (BRI). Full details of subtests and score construction are provided in the Supplementary Methods

### Statistical analysis

Differences across BMU-assigned profile groups were tested using one-way ANOVA for continuous variables and chi-square tests for categorical variables. Follow-up pairwise comparisons used Tukey HSD for ANOVA models, with Benjamini-Hochberg false discovery rate (BH-FDR) correction applied within families of related tests. For chi-square tests, standardised residuals were inspected to characterise over-and under-representation across cells. Profiles were analysed both categorically, using BMU island membership, and continuously, using centroid-distance scores. Sex and referral-status differences in centroid-distance scores were tested using Welch’s two-sample t-tests, and diagnosis-domain differences using one-way ANOVA with Tukey HSD post hoc tests. Cognitive outcomes and BRIEF T-scores were compared across BMU-assigned profile groups using one-way ANOVA with Tukey HSD post hoc tests. Effect sizes are reported as η^2^, Cohen’s d, and Cramér’s V. All tests were two-tailed with α=.05.

### Neural correlates of social profiles

In the MRI subsample, we constructed individual morphometric inverse divergence (MIND) networks^26^ from T1-weighted scans and tested multivariate covariation between neural morphologyand social profile expression. Structural MRI data were acquired at the CALM site using a 3T Siemens Prisma scanner with a 32-channel head coil^21^. T1-weighted images were collected using an MPRAGE sequence and pre-processed using the FreeSurfer recon-all surface-based pipeline (v7.4.0).

MIND networks^26^ were then constructed to quantify inter-regional structural similarity. The cortex was parcellated into 360 regions using the HCP-MMP1 atlas^27^, and five morphometric features were extracted for each region: cortical thickness, mean curvature, grey matter volume, sulcal depth, and surface area. These features were used to compute a 360 × 360 weighted similarity matrix for each participant, from which nodal strength was calculated for each parcel. Additional details are provided in the Supplementary Materials.

### Principal components analysis (PCA) and partial least squares (PLS)

To reduce the dimensionality of neuroimaging-derived regional strength values from the MIND networks, we ran a principal components analysis (PCA)^28^ using scikit-learn^29^. The number of retained principal components (PCs) was determined by parallel analysis based on 1,000 simulations of standardised random data. Components were retained when observed eigenvalues exceeded the 95% confidence interval of the simulated eigenvalue distribution. This indicated retention of 15 PCs, which together explained 20.4% of the variance in the original data (Figure 4A).

To determine the relationship between social profile expression and neuroimaging-derived structural similarity components, we ran a partial least squares (PLS) regression, with PCA-derived neuroimaging component scores entered as X and social profiles as Y. The number of latent variables was assessed using 10,000 permutations^30^ of Y. Component significance was evaluated by refitting the model and recomputing the partial correlation between X and Y component scores for each latent variable while controlling for age (at assessment), age at scan, and sex^31^. Components were retained when the observed partial correlation exceeded the null distribution generated by permutation. For the present data, this yielded a single significant component (p=0.02). Loading stability was assessed using 10,000 bootstrap resamples. Bootstrapped PLS loadings were aligned to the original solution using Orthogonal Procrustes rotation, and bootstrap ratios were calculated as the original X loadings divided by the standard deviation of the bootstrapped X loadings. Bootstrap ratios of ±2 were interpreted as stable loadings^32,33^.

### Software and data availability

Analyses were conducted in Python (v3.11.13) and R (v4.3.3). Python was used for data preparation, SOM modelling, neuroimaging, and statistical analyses, and R for network analyses. The code for analyses can be found here: (<u>GitHub</u>). CALM data are available via the CALM data portal, subject to the cohort’s data access procedures.

## Results

### Social dimensions from questionnaire items

Item-level relationships were modelled using a regularised partial correlation network, and communities were identified using Louvain modularity optimisation to derive core social dimensions (modularity Q=0.382; Figure S1, Text S1). Based on item content, the retained dimensions captured: (1) friendship behaviours (e.g. “has trouble keeping friends”); (2) social discomfort (e.g. “appears anxious in the company of other children”); (3) peer adversity (rejection and bullying) (e.g. “picked on or bullied by other children”); and (4) prosocial behaviour (e.g. “shares readily with other children”). These four dimensions formed the basis for constructing social profiles using a self-organising map.

### Self-organising map organisation of social profiles

The SOM arranged children with similar patterns across the four social dimensions into nearby locations on a two-dimensional grid, capturing graded variation across the sample. Map quality metrics indicated good representation of the input space (QE = 0.139; TE = 0.075; Text S2). Each SOM node stores a four-value weight vector, with one value corresponding to each of the four social dimension scores used to train the map. Component planes plot these values one dimension at a time, illustrating how the map represents variation in each dimension across the grid (Figure 2A).

The classification procedure identified significant SOM islands for all four archetypal profiles: social engagement, friendship difficulties, social withdrawal, and peer victimisation (Figure 2B). Island centroids were used to derive both categorical profile membership and continuous centroid-distance scores for each child, with children whose best-matching unit fell outside all significant islands classified as unassigned (see Methods for details Mean scores across the four social dimensions differed significantly by BMU-assigned profile (all p_FDR<1×10□ □ □), with each profile showing a distinct signature. Per-item mean scores across the original questionnaire items showed coherent item-level patterns consistent with the profile labels (Figure 2C). Together, this SOM-based classification approach translated graded variation across four social dimensions into interpretable, spatially localised profile regions, capturing non-linear combinations of behaviours that are not well summarised by any single dimension alone.

### Characterising social profiles

We next describe demographic and referral characteristics across SOM-derived social profiles (Table S1). Children were distributed across profiles as follows: friendship difficulties (n=15), social engagement (n=297), peer victimisation (n=31), social withdrawal (n=155), and unassigned (n=494). Children in the unassigned group had BMUs outside significant archetype islands and showed broadly intermediate scores, consistent with the null (median) reference profile.

Age did not differ across profiles (one-way ANOVA; q=.084). Sex distribution differed by categorical profile membership, with boys predominating across profiles and the highest proportion of girls in social engagement, χ^2^(4,N=992)=19.05, p<.001. Continuous centroid-distance analyses showed that boys were closer to the friendship difficulties, social withdrawal, and peer victimisation centroids, whereas girls were closer to the social engagement centroid (t-tests, q<.05; largest effect for social withdrawal, t(672.92)=-4.76, p<.001). Put simply, girls showed stronger social engagement, whereas boys showed stronger expression of the problem-oriented profiles.

Referral status also differed by categorical profile membership, with referral rates lowest in the social engagement profile, χ^2^(4,N=992)=47.79, p<.001. Continuous analyses showed that referred children were closer to the friendship difficulties, social withdrawal, and peer victimisation centroids, whereas non-referred children were closer to the social engagement centroid (Welch’s t-tests, all BH-FDR q≤3.81×10□□; largest effect for peer victimisation, t(344.20)=-15.63, p<.001). Overall, non-referred children showed stronger social engagement, whereas referred children showed stronger expression of the problem-oriented profiles (Figure 3A).

**Figure 3.**
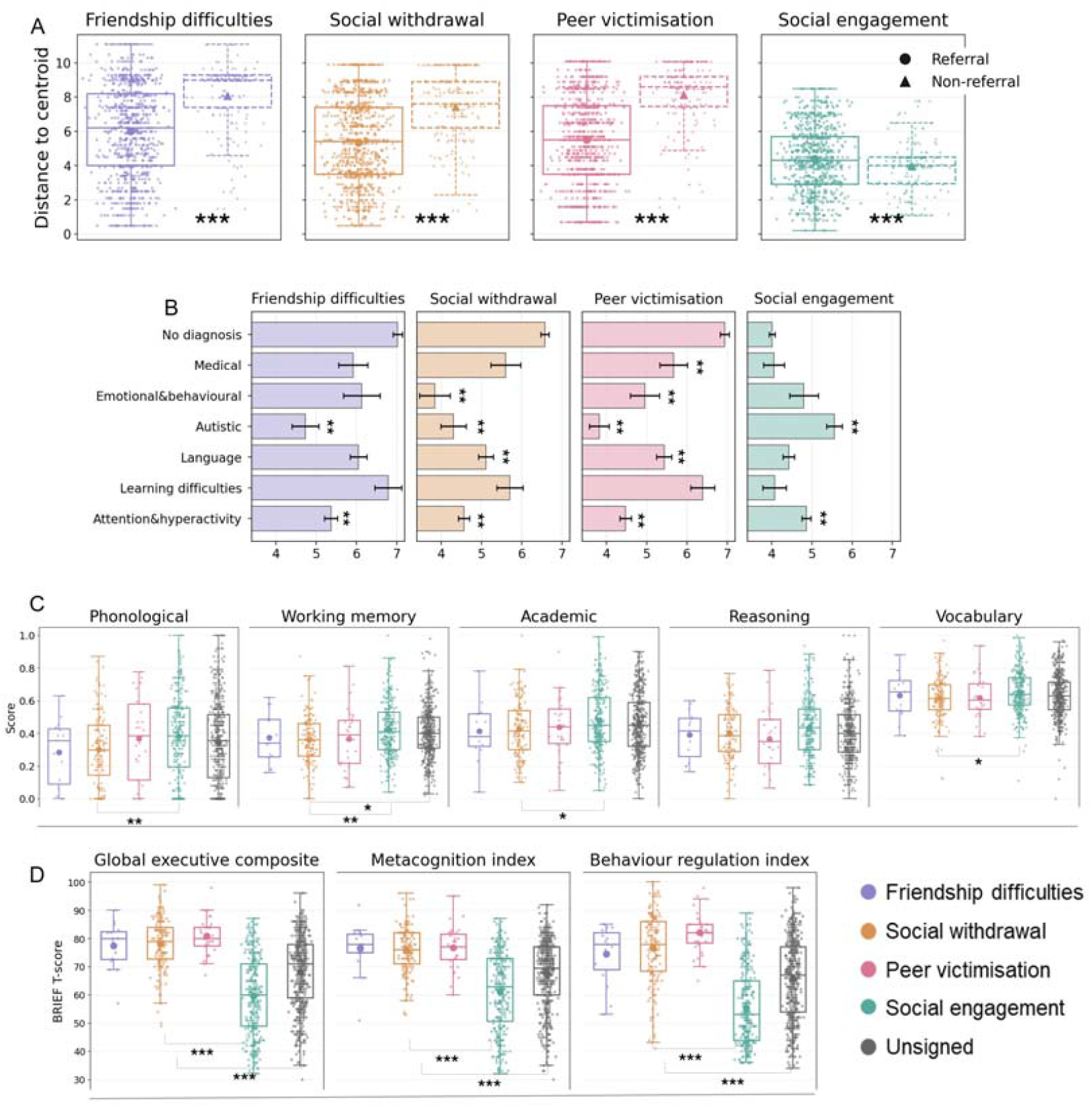
Referral, diagnosis, cognition and executive function across SOM-derived social profiles. (A) Centroid-distance scores by referral status (referred vs non-referred; lower values indicate greater proximity to each profile centroid), Welch’s t-tests with Benjamini–Hochberg FDR across centroids. (B) Mean centroid-distance to each profile centroid by diagnosis domain (mean ± SEM). Lower centroid-distance indicates greater proximity to the profile centroid, whereas higher centroid-distance indicates greater distance. For example, a higher distance to the social engagement centroid indicates a lower expression of the social engagement profile. (C) Cognitive scores across profiles (phonological processing, working memory, academic attainment, reasoning, vocabulary), one-way ANOVA with Tukey-adjusted post hoc tests. (D) BRIEF T-scores across profiles (global executive composite, metacognition index, behaviour regulation index; higher scores indicate greater difficulties), one-way ANOVA with Tukey-adjusted post hoc tests. Stars denote adjusted p values for the pairwise contrasts shown (post hoc for ANOVA panels): * < .05, ** < .01, *** < .001.

We next examined diagnosis-domain associations with continuous profile expression using centroid-distance scores (Figure 3B), where lower values indicate greater proximity to the profile centroid. Post hoc comparisons were made relative to children with no diagnosis, with mean differences reported as no diagnosis minus diagnosis (positive values indicate shorter distances in the diagnosis group). Attention/hyperactivity and autistic social showed shorter distances to friendship difficulties (1.70 and 2.49), social withdrawal (1.93 and 2.04), and peer victimisation (2.40 and 3.22), and longer distances to social engagement (−0.69 and −1.53; all p<.01). Language/communication and emotional/behavioural also differed from no diagnosis for social withdrawal (1.44 and 2.92) and peer victimisation (1.31 and 2.02), and neurological/medical differed for peer victimisation (1.27; all p<.01). Together, these patterns indicate greater proximity to difficulty-oriented profiles and reduced proximity to social engagement among children with attention/hyperactivity and autistic social diagnoses.

Cognitive performance differed by profile across five harmonised cognitive measures spanning phonological processing, working memory, academic attainment, reasoning, and vocabulary (Figure 3B). One-way ANOVAs showed profile effects for working memory, F(4,983)=4.42, p<.01, η^2^=.02, phonological processing, F(4,966)=3.00, p=.02, η^2^=.01, academic attainment, F(4,983)=2.61, p=.03, η^2^=.01, and weaker evidence for vocabulary, F(4,978)=2.39, p=.05, η^2^=.01, but not reasoning, F(4,985)=1.90, p=.11, η^2^=.01. Tukey-adjusted post hoc tests indicated higher scores in the social engagement profile than social withdrawal for academic attainment (mean difference=0.05, 95% CI [0.00, 0.10], p=.03), phonological processing (0.08, [0.01, 0.15], p=.01), vocabulary (0.04, [0.00, 0.07], p=.03), and working memory (0.06, [0.02, 0.10], p<.01). Working memory was also higher in the unassigned group than social withdrawal (0.04, [0.00, 0.08], p=.02). Overall, effects were driven mainly by lower cognitive scores in the social withdrawal profile compared with social engagement across multiple domains.

Executive function difficulties, indexed by BRIEF T-scores, differed across profiles for the behaviour regulation index, F(4,975)=80.22, p<.001, η^2^=.25, global executive composite, F(4,960)=68.55, p<.001, η^2^=.22, and metacognition index, F(4,961)=45.35, p<.001, η^2^=.16 (Figure 3C). Tukey-adjusted post hoc tests indicated lower BRIEF T-scores (fewer difficulties) in the social engagement profile than social withdrawal for the global executive composite (mean difference=18.46, 95% CI [15.07, 21.86], p<.001), metacognition (14.66, [11.39, 17.94], p<.001), and behaviour regulation (21.17, [17.51, 24.83], p<.001). Social engagement was also lower than the unassigned group across outcomes (global executive composite: 8.45, [5.93, 10.97], p<.001; metacognition: 6.34, [3.91, 8.77], p<.001; behaviour regulation: 10.26, [7.53, 12.98], p<.001). Together, these effects were driven primarily by consistently lower BRIEF T-scores in the social engagement profile compared with all other profiles, including the unassigned group.

### Neural correlates of social profiles

MIND networks^26^ quantify structural similarity across the cortex (Figure 4A). Regional strength values ranged from 11.93 to 65.21 (Figure 4B; M=43.15, SD=7.37). To reduce dimensionality and avoid testing 360 correlated regional strength features directly, we ran a PCA on strength across 360 regions. The number of components to retain was determined using parallel analysis (1,000 simulations of standardised random data), retaining components whose observed eigenvalues exceeded the 95% confidence interval of eigenvalues from the simulated datasets. This analysis indicated that 15 principal components (PCs) should be retained, which together explained 20.4% of the variance in the original strength data (Figure 4B). We then used PLS to relate strength PCs (X) to continuous social profile expression indexed by centroid-distance scores (Y), where lower distances indicate greater similarity to each profile centroid. Permutation testing identified one significant latent variable (p=0.02), capturing 4% of the covariance between MIND strength components and social profile expression (Figure 4C). Bootstrap ratios indicated that PCs 8, 11, 13 and 14 contributed stably to the latent brain pattern (Figure 4C). Using the loadings from these four PCs, we can visualise which regions are contributing most to the brain-social profile relationship in the PLS (Figure 4C). Regions within the top 10% of PC loadings were distributed across the cortex, with the largest concentration in frontal (12 regions) and parietal (15 regions) areas, followed by occipital (4 regions) and temporal (4 regions) lobes. Contributing regions were broadly balanced across hemispheres, with slightly more in the left hemisphere (20 left, 16 right) (Figure 4C). Y-loadings indicated that the brain-behaviour covariance was most strongly expressed for social withdrawal and peer victimisation, with an opposite loading for social engagement (Figure 4C).

**Figure 4.**
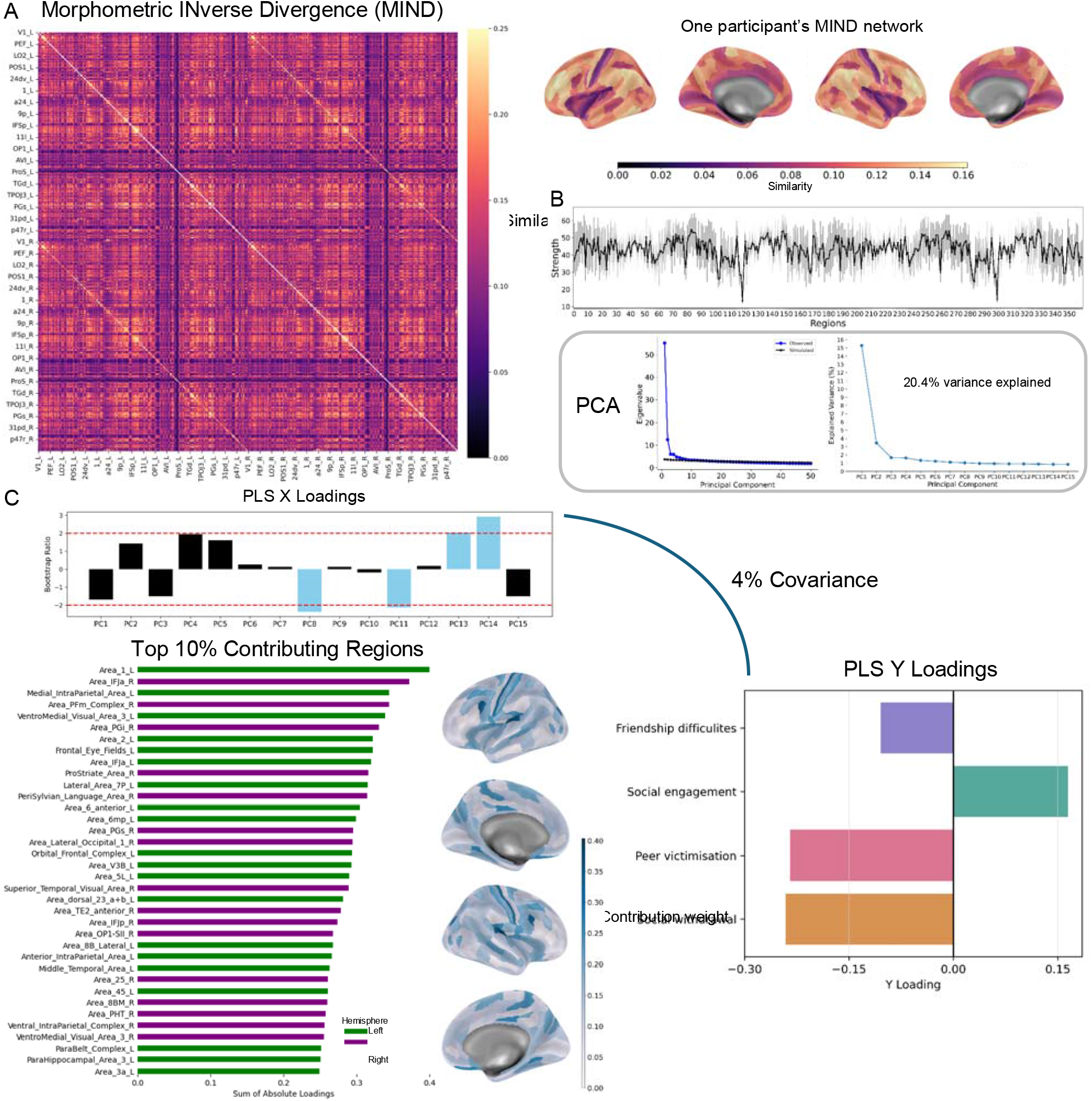
MIND networks and their relationship with social profile expression. (A) Group-average MIND similarity matrix (left) and an illustrative participant’s MIND network projected onto the cortical surface (right). (B) Regional strength across the 360 cortical regions, and PCA of regional strength. The retained 15 principal components explained 20.4% of variance in the original regional strength data. (C) PLS relating strength PCs (X) to centroid-distance social profile expression scores (Y). One significant latent variable (p = 0.019) captured 4% covariance between brain and social measures. Bootstrapped ratios of X loadings indicate stable contributions from PCs 8, 11, 13 and 14. The surface plots show the sum of absolute loading of each region to these four PCs. The bar plot shows the top 10% contributing regions based on absolute loadings summed across these stable PCs, indicating broad distribution across hemispheres with a predominance of frontal and parietal regions. Y-loadings indicate that the covariance pattern is driven primarily by social withdrawal and peer victimisation, with an opposite loading for social engagement.

## Discussion

In transdiagnostic neurodevelopmental research, social functioning is often treated as secondary, despite its central role in children’s everyday adaptation and learning. Here, we treat social behaviour and peer experiences as a primary phenotype and show that they are not well captured by a single dimension. Instead, distinct profiles of co-occurring social behaviours and peer adversity were systematically associated with referral and diagnosis, cognitive performance, and T1-derived morphometric network organisation.

Foregrounding social behaviour and peer experiences is warranted because these dimensions are tightly linked to children’s learning opportunities and adjustment. Meta-analytic evidence indicates that peer social acceptance is positively associated with academic achievement^8^, whereas peer victimisation is associated with lower academic achievement^5^ and with reductions in cognitive-motivational resources that support learning^34^. Peer adversity is also linked to elevated risk for later mental health difficulties and broader life outcomes^4^. Importantly, the peer context is a feasible intervention target: universal school-based social and emotional learning programmes improve academic performance^19^, and anti-bullying programmes reduce bullying and victimisation^35^.

Some children struggle to form and maintain friendships, others withdraw, and others are exposed to victimisation, while some show relatively high prosocial engagement. We therefore moved beyond single-scale summaries by using an item-level, data-driven approach to map graded profiles of co-occurring social behaviours and peer experiences in a large transdiagnostic cohort.

We identified four distinct profiles: social engagement, friendship difficulties, social withdrawal, and peer victimisation, with some children falling outside the profile islands and showing more mixed, intermediate patterns. We also derived centroid-distance scores indexing graded profile expression.

These social signatures were clinically meaningful, tracking systematic differences by sex, referral, and diagnosis. Boys showed greater proximity to the more problem-oriented profiles, whereas girls were closer to social engagement. Referral was least common in social engagement. Attention/hyperactivity and autistic social diagnoses showed the strongest shift towards friendship difficulties, withdrawal, and victimisation, alongside reduced proximity to social engagement. Other domains showed smaller shifts, with neurological/medical and learning difficulties differing little from the no-diagnosis group. Taken together, social difficulties were transdiagnostic, but concentrated most strongly within specific neurodevelopmental pathways captured by referral and diagnosis, highlighting the value of profiling social risk signatures beyond diagnostic categories.

The distinct social profiles were also linked to broader functional outcomes. Cognitive differences were most apparent for social withdrawal, which showed lower performance than social engagement across working memory, phonological processing, academic attainment, and vocabulary. In contrast, reasoning showed no clear profile differentiation, suggesting that social profile variation aligned more with learning- and language-related skills than with non-verbal reasoning. Everyday executive function showed a different pattern: children in the social engagement profile had fewer difficulties across BRIEF indices than all other profiles, including the unassigned group. Together, these findings suggest that social difficulties are not uniformly linked to wider difficulties, but may map onto distinct patterns of cognitive and self-regulatory strengths and vulnerabilities.

A central implication of these findings is that children’s social behaviour should not be viewed solely as a secondary consequence of diagnosis or cognitive difficulty, but as a meaningful developmental context that may shape downstream cognitive and regulatory outcomes. Much of the literature has characterised peer problems as a common feature of neurodevelopmental conditions such as ADHD, with downstream consequences for adjustment and learning^36^. In contrast, our results show that variation in social engagement is systematically associated with cognitive performance, everyday executive functioning, and brain structure, consistent with the view that social behaviour may contribute to developmental cascades rather than sitting at their periphery.

Notably, children in the social engagement profile showed the most consistently positive pattern of functioning across outcomes. They showed fewer everyday executive function difficulties and tended to show higher cognitive performance than other profiles, particularly social withdrawal, and in some cases even relative to the unassigned group.

These findings are cross-sectional and do not establish directionality. However, longitudinal and meta-analytic evidence indicates that stronger peer relationships and social competence are prospectively associated with better academic outcomes^8^. Complementing this, longitudinal work also links prosocial behaviour with later executive function during early childhood^37^. Together, this evidence is consistent with the hypothesis that social engagement may support cognitive and self-regulatory development over time, while cognitive self-regulation may also support successful peer engagement, motivating longitudinal and mechanistic intervention studies to test directionality and pathways.

Structural neuroimaging findings also suggest that social engagement is not only behaviourally distinct, but associated with differences in brain structure. In the morphometric network analysis, social profile expression covaried with structural network patterns, with social engagement showing loadings in the opposite direction to the more problem-oriented profiles. Although the explained variance was modest and the data are cross-sectional, this dissociation suggests that social engagement is embedded within broader neurodevelopmental organisation, rather than reflecting a purely descriptive behavioural pattern. This aligns with prior work linking prosocial behaviour and peer experiences to individual differences in cortical development and broader structural and functional brain outcomes^38–40^.

Together, these findings invite a shift in perspective. Rather than asking only why children with cognitive or neurodevelopmental difficulties struggle socially, it may be equally important to ask how children’s social environments and behaviours contribute to cognitive and self-regulatory development over time. Social engagement may provide richer opportunities for learning, language use, and practice of regulatory skills, whereas withdrawal or peer adversity may constrain these opportunities. Our results suggest that treating social behaviour as a primary developmental exposure may be critical for understanding heterogeneity in cognitive and executive outcomes across childhood.

Several limitations should be noted. First, the analyses are cross-sectional, so they cannot adjudicate directionality between social profiles, cognition, and brain structure. Cross-sectional associations can also misrepresent longitudinal processes, particularly when interpreting cascade-like pathways^41^. Second, social functioning was indexed using parent report only. Parent reports capture observable behaviours but do not directly assess children’s subjective experiences, including loneliness, and informants often show meaningful discrepancies in child assessment^42^. Third, two profiles were small in absolute size, particularly friendship difficulties and peer victimisation, reducing precision for profile-specific estimates and limiting the scope for subgroup analyses. Finally, neuroimaging analyses were conducted in an MRI subsample, which reduces power and may introduce selection effects if scan availability and data quality relate to behaviour or clinical factors.

Future work should test directionality using longitudinal designs with repeated assessments of social signatures, cognition and executive function, and should incorporate multi-informant and school-context measures to triangulate parent report. Intervention studies can then test whether strengthening peer contexts and prosocial engagement shifts cognitive and regulatory trajectories over time.

### Conclusion

In a large transdiagnostic neurodevelopmental cohort, we show that social functioning is not unitary, but organises into graded profiles of social engagement, friendship difficulties, social withdrawal, and peer victimisation, alongside a substantial intermediate group. These social signatures were clinically informative, tracking systematic differences in sex, referral, and diagnostic pathways. They were also functionally meaningful: social withdrawal showed the clearest disadvantage across learning- and language-related cognitive domains, whereas social engagement was associated with markedly fewer everyday executive function difficulties across BRIEF indices. Finally, social profile expression covaried with T1-derived morphometric network organisation, indicating that social signatures align with broader neurodevelopmental organisation. Together, these findings position social behaviour and peer experiences as a primary developmental phenotype, motivating longitudinal and intervention studies to test directionality and to determine whether shifting children’s social signatures can improve cognitive and self-regulatory trajectories.

## Acknowledgements

The authors thank the children and families who participated in the study. The authors also thank Kayson Fakhar, PhD, and William Mills, BA, for their support and readiness to help throughout the project.

