## Supplementary materials for "Mapping social profiles in childhood and adolescence: associations with cognition and brain structure"

**Supplementary Methods**

*Diagnoses information*

Diagnostic information was collated from referrer-supplied clinical history at recruitment, consistent with the CALM protocol ^1^. Children in CALM were referred by education or health professionals for difficulties with attention, learning, and/or memory, irrespective of diagnostic status. Of the full sample, 805 children were referred, and 187 were non-referred. Non-referred children were recruited from the same schools and communities and served as a community comparison group within the CALM cohort. Referrals were primarily from education services (63%), with additional referrals from clinical services (33%) and speech and language therapists (4%) ^2^. For analysis, diagnostic indicators were grouped into broad domains reflecting common referral and diagnostic pathways in CALM: attention/hyperactivity (e.g., ADHD, probable ADHD), autistic social (e.g., autism, PDA), language/communication (e.g., language disorder, speech and language therapy support), learning difficulties (e.g., dyslexia, dyscalculia, dyspraxia), emotional/behavioural (e.g., anxiety, OCD, depression, conduct disorder, ODD), and neurological/medical (e.g., epilepsy, known genetic condition, FASD). Children without any recorded diagnosis were coded as ‘no diagnosis’ (Table S1).

*Self-organising map (SOM)*

The SOM was trained on each child’s four social dimension scores, scaled from 0 to 1, using a 10 × 10 rectangular grid with Euclidean activation distance. Models were fit using MiniSom (Vettigli, 2013/2026). Training was performed for 5,000 iterations with random weight initialisation, a Gaussian neighbourhood function (sigma = 2), and learning rate 0.5. Each participant was mapped to their best matching unit (BMU), defined as the node whose weight vector was closest to the participant’s input vector (Kohonen, 2001). Map quality was summarised using quantisation error and topographic error (QE = 0.1389, TE = 0.0746).

To identify interpretable regions within the SOM, we used a supervised classification approach anchored by simulated archetype profiles. Four archetypes were generated to reflect social engagement, friendship difficulties, social withdrawal, and peer victimisation. For each archetype, the corresponding dimension was set to the 95th percentile and the remaining dimensions to the 50th percentile, using the same 0 to 1 scaling as the empirical data. A null archetype reflecting median values across all dimensions was included as a reference. For each archetype, 500 simulated observations were generated.

Archetype-associated regions were identified using one-vs-rest support vector machine (SVM) classifiers with a radial basis function kernel (Cortes & Vapnik, 1995). Classifiers were trained on the simulated archetype observations, with balanced class weights to address class imbalance, and then applied to SOM neuron weight vectors. Each neuron was assigned to the archetype with the highest decision score.

We then tested whether archetype-labelled regions formed non-random spatial islands on the SOM using a permutation-based island-size procedure. Islands were defined as connected components of same-labelled neurons using 8-neighbour connectivity. Null distributions of island sizes were generated using 1,000 permutations in which SOM neuron weight vectors were randomly reassigned to grid locations, reclassified using the same one-vs-rest SVM procedure, and island sizes recomputed. For each archetype, the minimum island size required for significance was defined as the 99th percentile of its null distribution (p < .01). Significant islands were retained, and island centroids were computed as the mean row and column coordinates of neurons within each island.

Profile outputs were derived in two ways. For categorical assignment, children were assigned to the profile island containing their BMU, and those whose BMU fell outside all significant islands were classified as unassigned. For continuous profile expression, Euclidean distance was computed between each child’s BMU coordinate and the centroid of each significant island, with lower values indicating greater proximity to, and stronger expression of, the corresponding profile. Each child therefore had both a categorical profile label and four continuous centroid-distance scores, one per profile (Figure 2B).

*Cognitive measures*

Cognitive performance was assessed using standardised measures of phonological processing, working memory, reasoning, vocabulary, and academic attainment. Phonological processing was measured using the Phonological Assessment Battery (PhAB) ^3^, reported as standard age scores (M = 100, SD = 15). Working memory was assessed using the Automated Working Memory Assessment (AWMA) ^4^, reported as standard scores (M = 100, SD = 15). Reasoning was indexed using the WASI-II ^5^, (Matrix Reasoning T-score, and where available FSIQ-2 or FSIQ-4, M = 100, SD = 15). Vocabulary was assessed using the Peabody Picture Vocabulary Test (PPVT-4 or PPVT-5) ^6^, reported as age-based standard scores (M = 100, SD = 15). Academic attainment was assessed using the WIAT-II subtests Spelling, Word Reading, and Numerical Operations ^7^, reported as standard scores (M = 100, SD = 15).

We derived five cognitive summary measures from age-standardised scores and rescaled each summary measure to a 0–1 range (higher values indicate better performance) for profile comparisons. Only working memory and academic attainment were averaged across subtests; the other three outcomes used a single standard score each. Two summary measures were composites, calculated as the participant-level mean of constituent standard scores. Working memory was the mean of four AWMA standard scores (Digit Recall, Dot Matrix, Backward Digit Recall, Mr X). Academic attainment was the mean of three WIAT-II standard scores (Spelling, Word Reading, Numerical Operations). The remaining three summary measures were single test scores analysed directly: phonological processing (PhAB Object Naming and rapid automatised naming combined standard score), reasoning (Matrix Reasoning T-score), and vocabulary (PPVT standard score). These five harmonised cognitive measures were used in all downstream profile comparisons.

*Executive function (BRIEF)*

Everyday executive function was assessed using the parent-report Behaviour Rating Inventory of Executive Function. The BRIEF comprises 80 items describing everyday behaviours over the past six months, rated on a three-point scale. Raw scores were converted to age- and sex-normed T-scores (M = 50, SD = 10), with higher scores indicating greater executive function difficulties. Analyses focused on the three composite indices: the Global Executive Composite (GEC), Metacognition Index (MI), and Behaviour Regulation Index (BRI) ^8^.

*Neural correlates of social profiles*

Neuroimaging analyses were conducted in a subsample of children with associated MRI data. Structural MRI data were acquired at the CALM site using a 3T Siemens Prisma scanner with a 32-channel head coil. T1-weighted images were collected using a magnetisation-prepared rapid gradient echo (MPRAGE) sequence. Full acquisition parameters are reported in the CALM protocol ^1^.

T1-weighted images were preprocessed using FreeSurfer v7.4.0 with the recon-all surface-based pipeline ^9^. Surface reconstruction quality was indexed using the Euler number, an automated proxy for topological defects, where lower values indicate poorer reconstruction quality. Two participants showed reduced Euler values when summed across hemispheres and were flagged during quality control.

For each participant, morphometric inverse divergence (MIND)^10^ networks were constructed to quantify inter-regional structural similarity . The cortex was parcellated into 360 regions using the HCP-MMP1 atlas ^11^. For each region, five morphometric features were extracted: cortical thickness, mean curvature, grey matter volume, sulcal depth, and surface area. These features were used to compute a 360 × 360 weighted similarity matrix for each participant, representing the MIND network.

To summarise each participant’s MIND network at the regional level, we computed nodal strength for each of the 360 parcels using the Brain Connectivity Toolbox (BCTPY) in Python 3.11.13 ^12^. In weighted networks, nodal strength is defined as the sum of the weights of all edges connecting a given node to every other node. This yielded a 360-element vector of regional strength values for each participant.

**Supplementary Results**

**Table S1**

|  | Friendship difficulties | Social engagement | Peer victimisation | Social withdrawal | Unassigned | Total |
| --- | --- | --- | --- | --- | --- | --- |
| N | 15 | 297 | 31 | 155 | 494 | 992 |
| Age, years mean (SD) | 9.19 (1.69) | 9.47 (2.10) | 9.93 (2.48) | 10.06 (2.72) | 9.50 (2.45) | 9.59 (2.39) |
| Girls n (%) | 4 (26.7) | 131 (44.1) | 8 (25.8) | 45 (29.0) | 151 (30.6) | 339 (34.2) |
| Boys n (%) | 11 (73.3) | 166 (55.9) | 23 (74.2) | 110 (71.0) | 343 (69.4) | 653 (65.8) |
| Referred n (%) | 15 (100.0) | 212 (71.4) | 30 (96.8) | 148 (95.5) | 400 (81.0) | 805 (81.1) |
| Attention/hyperactivity | 6 (40.0) | 42 (14.1) | 15 (48.4) | 66 (42.6) | 130 (26.3) | 259 (26.1) |
| Autistic social | 3 (20.0) | 2 (0.7) | 6 (19.4) | 20 (12.9) | 31 (6.3) | 62 (6.2) |
| Language/communication | 3 (20.0) | 44 (14.8) | 5 (16.1) | 34 (21.9) | 81 (16.4) | 167 (16.8) |
| Learning difficulties | 0 (0.0) | 15 (5.1) | 0 (0.0) | 9 (5.8) | 27 (5.5) | 51 (5.1) |
| Emotional/behavioural | 1 (6.7) | 6 (2.0) | 1 (3.2) | 12 (7.7) | 13 (2.6) | 33 (3.3) |
| Neurological/medical | 1 (6.7) | 19 (6.4) | 3 (9.7) | 8 (5.2) | 18 (3.6) | 49 (4.9) |
| No diagnosis | 5 (33.3) | 192 (64.6) | 9 (29.0) | 44 (28.4) | 257 (52.0) | 507 (51.1) |

***Table S2. Social items included in the network analysis (22 items)***

| Measure | Item number | Item text |
| --- | --- | --- |
| SDQ peer problems | 6 | Rather solitary, tends to play alone. |
| SDQ peer problems | 11 | Has at least one good friend. |
| SDQ peer problems | 14 | Generally liked by other children. |
| SDQ peer problems | 19 | Picked on or bullied by other children. |
| SDQ peer problems | 23 | Gets on better with adults than with other children. |
| SDQ prosocial | 1 | Considerate of other people’s feelings. |
| SDQ prosocial | 4 | Shares readily with other children. |
| SDQ prosocial | 9 | Helpful if someone is hurt, upset or feeling ill. |
| SDQ prosocial | 17 | Kind to younger children. |
| SDQ prosocial | 20 | Often volunteers to help others. |
| CCC-2 social | 3 | Appears anxious in the company of other children. |
| CCC-2 social | 7 | With familiar adults, seems inattentive, distant, or occupied. |
| CCC-2 social | 13 | Teased or bullied by other children. |
| CCC-2 social | 16 | Left out of joint activities by other children. |
| CCC-2 social | 33 | Hurts or upsets other children without meaning to. |
| CCC-2 social | 57 | Shows concern when other people are upset. |
| CCC-2 social | 67 | Talks about friends and shows interest in what they do and say. |
| Conners-3 peer relations | 4 | Is one of the last to be picked for games. |
| Conners-3 peer relations | 6 | Does not know how to make friends. |
| Conners-3 peer relations | 18 | Has trouble keeping friends. |
| Conners-3 peer relations | 38 | Has no friends. |
| Conners-3 peer relations | 43 | Does not get invited to play or go out with others. |

**Figure S1. Louvain community structure of the social-item network**. Nodes represent individual questionnaire items from the SDQ, CCC-2, and Conners-3 (see Table S2), and edges indicate regularised partial correlations. Node colours denote Louvain communities (scaled edge weights).

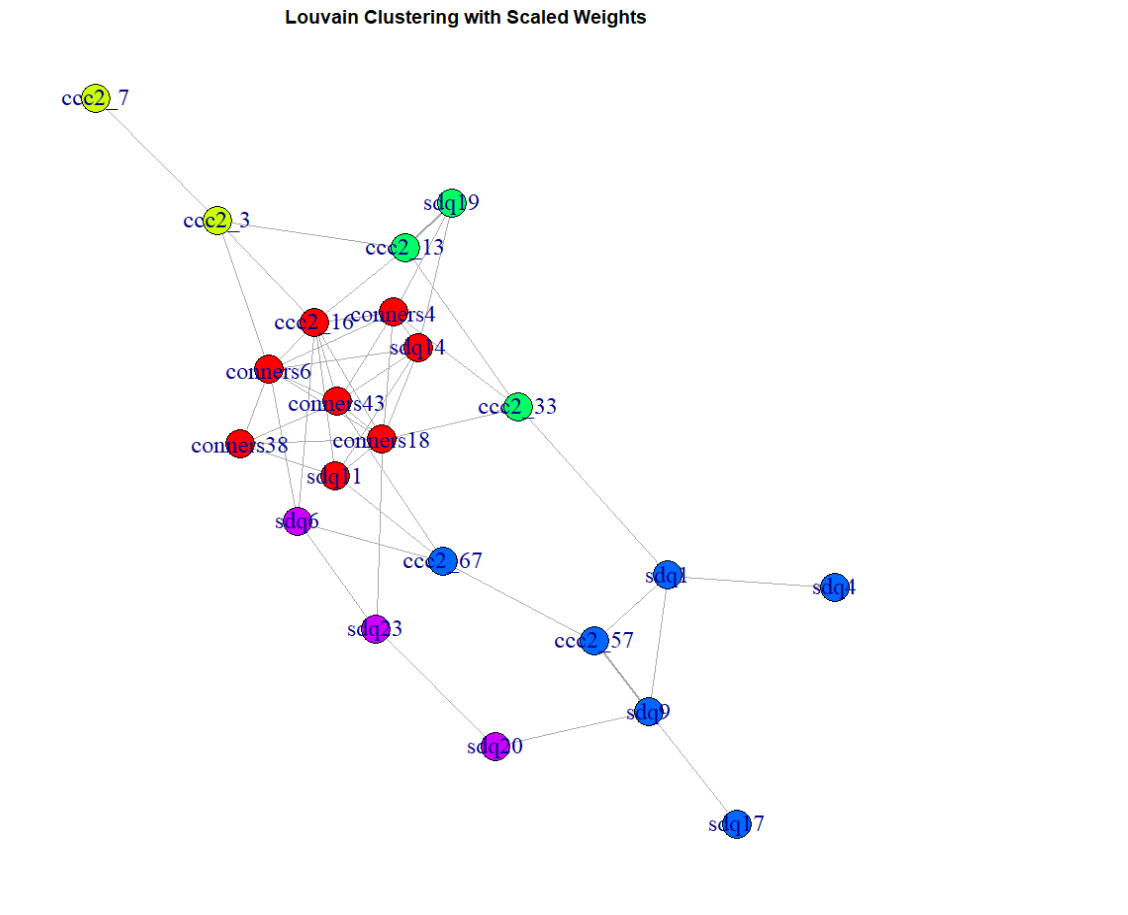

***Text S1 Social dimensions from questionnaire items***

Item-level relationships were modelled using a regularised partial correlation network, and communities were identified using Louvain modularity optimisation to derive core social dimensions (modularity Q=0.382; Figure S1). Four dimensions showed acceptable-to-excellent reliability (α=0.924, 0.545, 0.761, 0.835). The dimension with α = 0.545 contained only two items and was retained for conceptual coherence, given the constrained item count. The fifth dimension showed poor internal consistency (α = 0.236) and was excluded, leaving four coherent dimensions comprising 19 items.

***Text S2. SOM diagnostics and clustering analyses***

Inspection of the SOM distance map (U-matrix) showed smooth transitions across the grid rather than sharply delineated boundaries, consistent with graded variation in social behaviour (Figure S2A). Best-matching unit (BMU) activation frequencies indicated that children were distributed across the map rather than concentrated within a small number of discrete regions (Figure S2B). Map quality indices are reported in the main text (QE = 0.139; TE = 0.075).

To assess whether the SOM could be partitioned into discrete subgroups, we applied standard clustering methods to the SOM representation. Silhouette support declined as the number of clusters increased, and visual inspection suggested that clustering solutions did not yield clearly separable or stable boundaries across the map (Figure S2C-D). These analyses supported the interpretation of graded variation across the SOM and informed the use of the archetype-anchored classification approach in the main analyses.

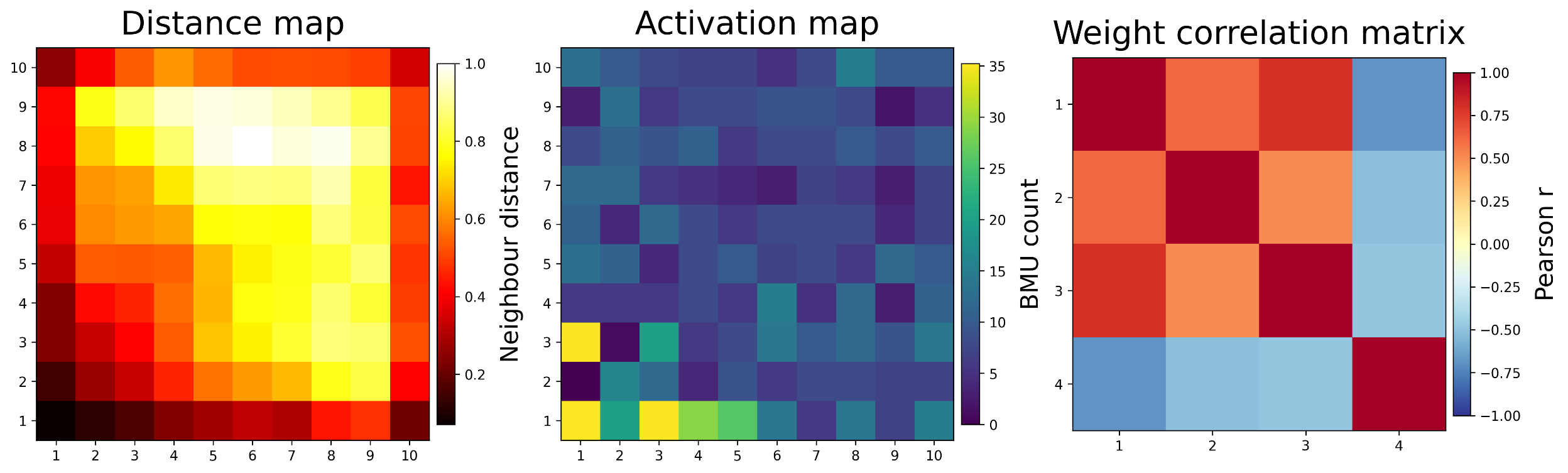

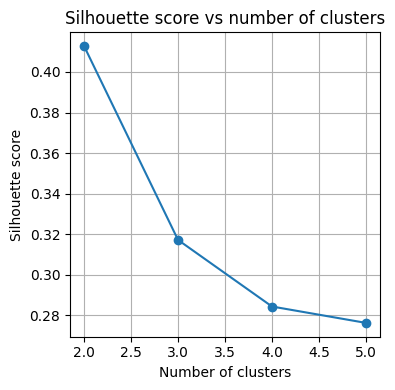

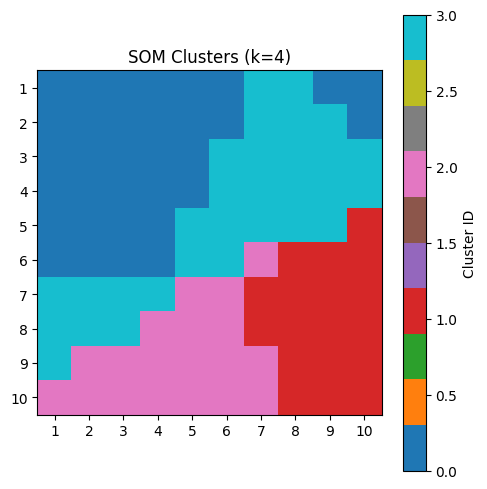

**Figure S2. SOM diagnostics and clustering support**. (A) U-matrix (distance map) showing neighbourhood distances across the 10 × 10 SOM grid. (B) Activation map showing best-matching unit (BMU) frequencies across the grid. (C) Silhouette scores across cluster solutions, indicating reduced support with increasing numbers of clusters. (D) SOM clustering solution for k = 4. Cluster assignment of SOM nodes across the 10 × 10 grid, illustrating that clusters form broad, contiguous regions rather than sharply separable subgroup boundaries.

A

C

D

B
